# Dissociable thalamic oscillatory mechanisms support motor sequence learning

**DOI:** 10.64898/2026.07.29.741410

**Authors:** Angela Voegtle, Lars Buentjen, Stefan Repplinger, Slawomir J. Nasuto, Adriano de Oliveira Andrade, Matthias Deliano, Robert T. Knight, Richard B. Ivry, Catherine M. Sweeney-Reed

**Affiliations:** Neurocybernetics and Rehabilitation, Dept. of Neurology, Otto von Guericke University Magdeburg, Magdeburg, Germany; Dept. of Stereotactic Neurosurgery, Otto von Guericke University Magdeburg, Magdeburg, Germany; Dept. of Neurology, Otto von Guericke University Magdeburg, Magdeburg, Germany; Biomedical Sciences and Biomedical Engineering Division, School of Biological Sciences, University of Reading, Reading, United Kingdom; Centre for Innovation and Technology Assessment in Health (NIATS), Federal University of Uberlândia (UFU), Brazil; Faculty of Electrical Engineering (FEELT), Postgraduate Program in Biomedical (PPGEB), Uberlândia, Brazil; Combinatorial Neuroimaging Core Facility, Leibniz Institute for Neurobiology, Magdeburg, Germany; Helen Wills Neuroscience Institute, University of California—Berkeley, Berkeley, California, USA; Depts. of Neuroscience and Psychology, University of California—Berkeley, Berkeley, California, USA; Center for Behavioral Brain Sciences - CBBS, Otto von Guericke University Magdeburg, Magdeburg, Germany

**Keywords:** motor sequence learning, intracranial recording, ventrointermediate nucleus of the thalamus, serial reaction time task, phase amplitude coupling

## Abstract

The ventrointermediate nucleus of the thalamus (VIM) is implicated in motor sequence learning, yet the underlying neural mechanisms remain unclear. We recorded intracranial activity from the human VIM during a serial reaction time task to determine how neural dynamics support learning. Participants responded faster during repeating than randomized sequences. Beta-to-low-gamma activity was greater during repeating sequences and elevated relative to the prestimulus baseline, consistent with emergence and stabilization of learned motor representations. In contrast, beta-phase modulation of high-frequency activity decreased progressively from rest to random to repeated sequence execution. Stronger phase–amplitude coupling was associated with faster responses, reaching significance in the random condition. These findings reveal a dissociation between power and cross-frequency coupling. Together they suggest that thalamic dynamics contribute to motor learning through multiple mechanisms: beta-band power reflects the emergence of learned motor representations, whereas beta–high-frequency coupling is enhanced when the context is less predictable.

## Introduction

The cortico–subcortical network underlying motor learning involves the cerebellum, basal ganglia, primary motor cortex, and the ventrointermediate nucleus of the thalamus (VIM)^1–3^. A functional network involving these structures is associated with tremor, with a central role for the VIM highlighted by the tremor reduction observed from VIM with deep brain stimulation (DBS)^4,5^. Here we used DBS implants to examine the contribution of the human VIM to motor learning^6^.

Beta oscillations within cortico–subcortical motor circuits are dynamically modulated during movement and learning^1,7–9^. Thalamic, basal ganglia, and cortical beta activity is typically suppressed during movement execution^1,7,10–12^, and the magnitude of this suppression is enhanced during unfamiliar or demanding behavior^13^. In familiar or well-learned behavioral contexts, increases in beta power are observed relative to unfamiliar contexts ^9,13^, with prominent beta band activity observed during the resting state^10^. These observations have led to the proposal that beta oscillations reflect the maintenance or stabilization of the current motor state (“status quo”)^10^, with enhanced beta observed not only during rest (stable motor state) but also during the production of learned movements^13^. Consistent with this interpretation, modulation of VIM activity through DBS influences both motor stability (e.g., reduced tremor) and sequence learning performance^6,8^.

It remains unclear how local oscillatory dynamics within the VIM contribute to the representation and execution of learned motor sequences. Cross-frequency coupling has been proposed as a mechanism for coordinating cortical– subcortical neural activity across spatial and temporal scales^14–16^. Phase–amplitude coupling (PAC), in which the amplitude of higher-frequency activity is modulated by the phase of a low-frequency oscillation, has been observed in cortical and subcortical structures across cognitive and motor tasks^14,17–20^. In motor systems, coupling of high-frequency activity (HFA) with beta activity is dynamically modulated during movement, typically decreasing around movement execution. Moreover, this coupling is pathologically elevated at rest in movement disorders such as Parkinson’s disease and, when attenuated, is associated with clinical improvement^19,21,22^. Although PAC has been observed within the thalamus and in thalamo-cortical interactions^15,23,24^, its role in motor sequence learning in the thalamus is unknown.

Here, we investigated how oscillatory activity and cross-frequency coupling within the VIM are modulated during motor sequence learning using intracranial recordings in patients with essential tremor performing a serial reaction time task (SRTT; Fig. 1a)^25^. We compared neural dynamics during sequential movements in which the successive responses either followed a repeating sequence or were determined randomly, asking how thalamic activity was modulated by the presence or absence of sequential structure.

**Fig. 1.**
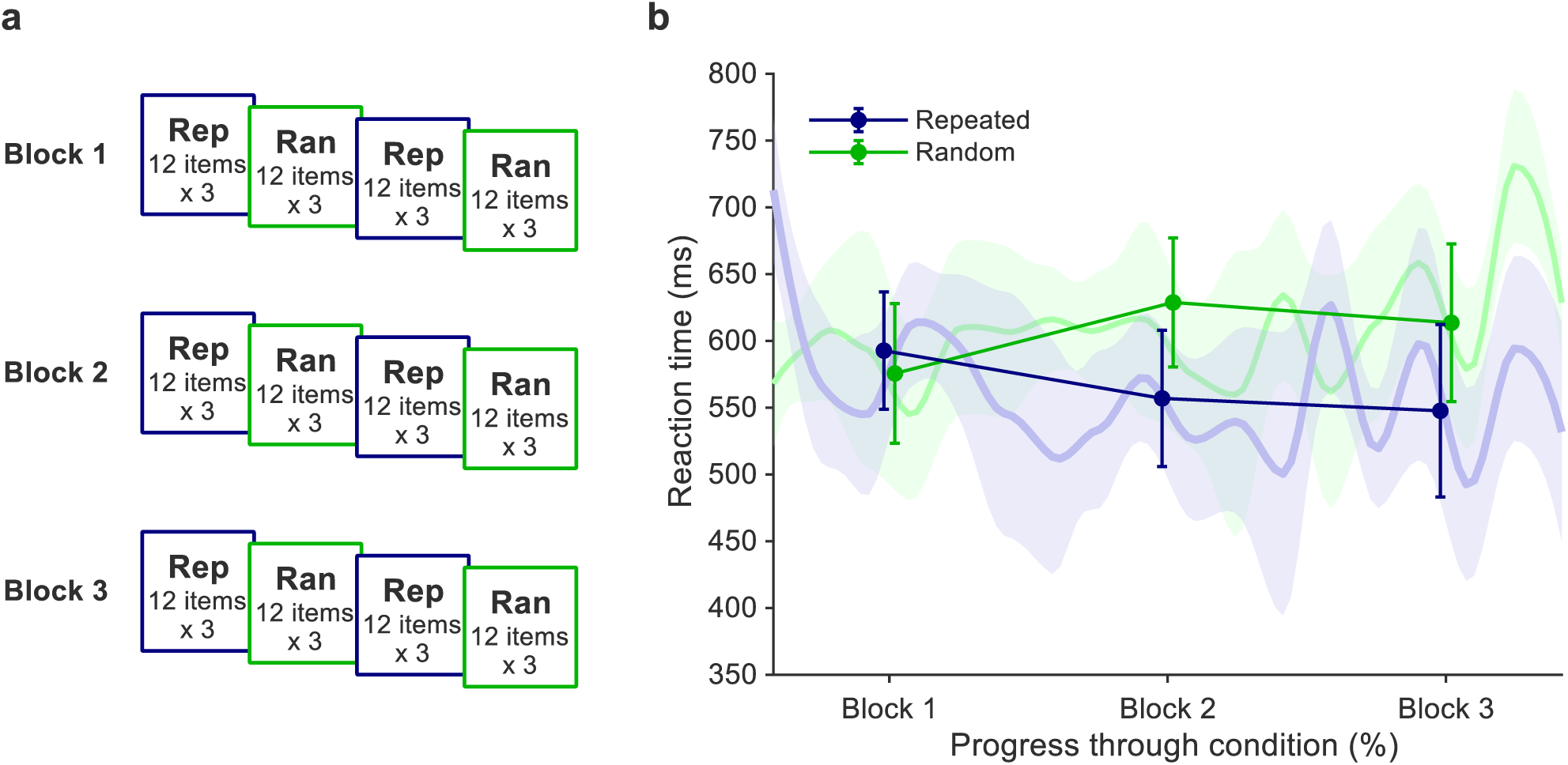
Behavioral experiment and performance. **a** Serial reaction time task, with alternating runs of repeated (Rep) sequences and random (Ran) trials across three blocks. **b** Reaction times (RTs) across Rep and Ran (mean ± s.e.m.). RTs were faster during Rep (F(1,6) = 24.8, p = 0.002, *η_p_*^2^ = 0.81).

## Results

### Behavioral performance

Reaction times (RTs) were faster during repeated sequences than random trials (main effect of *Sequence type*: F(1,6) = 24.8, p = 0.002, η*_p_*^2^ = 0.81; Fig. 1b) indicating that the participants exhibit significant motor learning during the experiment. There was no main effect of *Time* (F(2,12) = 0.16, p = 0.86, η*_p_*^2^ = 0.026), and the *Sequence type* × *Time* interaction only approached significance (F(2,12) = 3.85, p = 0.051, η*_p_*^2^ = 0.39). Taken together, these results indicate that reaction time remained relatively stable across the three blocks despite learning during the repeated blocks. This may reflect a balance between learning-related improvements and other factors such as fatigue or ceiling effects. Post hoc comparisons were performed to assess the direction of the effects, given the interaction was close to significant (p = 0.051). Post hoc comparisons showed that RTs were faster during Repeated than Random in Blocks 2 and 3 (p = 0.023 and p = 0.015, respectively). Accuracy did not differ between conditions or across blocks (main effect of *Sequence type*: F(1,6) = 1.09, p = 0.34, η*_p_*^2^ = 0.15; main effect of *Time*: F(2,12) = 0.50, p = 0.62, η*_p_*^2^ = 0.077; *Sequence type* × *Time* interaction: F(2,12) = 0.16, p = 0.86, η*_p_*2 = 0.025). Accuracy was slightly greater for Repeated than Random trials in each block, precluding a speed–accuracy trade-off.

### Learning-related modulation of oscillatory activity

Event-related spectral perturbations (ERSPs) showed greater beta–low-gamma power during Repeated than Random trials. Cluster-based permutation testing confirmed a significant difference between conditions from approximately 500 to 800 ms after stimulus onset in a window spanning 13–41 Hz (p_pos_ = 0.023, paired Cohen’s d_z_ = 0.65), Fig. 2a-c). Post hoc examination of cluster-averaged values indicated that, relative to a prestimulus baseline, power within the cluster was significantly greater during Repeated trials (mean ± s.e.m.: 1.96 ± 0.69 dB; p = 0.039). In contrast, power during Random trials did not differ significantly from this baseline (−1.51 ± 1.39 dB; p = 0.55). This pattern indicates that sequence learning was associated with the emergence of beta–low-gamma activity.

**Fig. 2.**
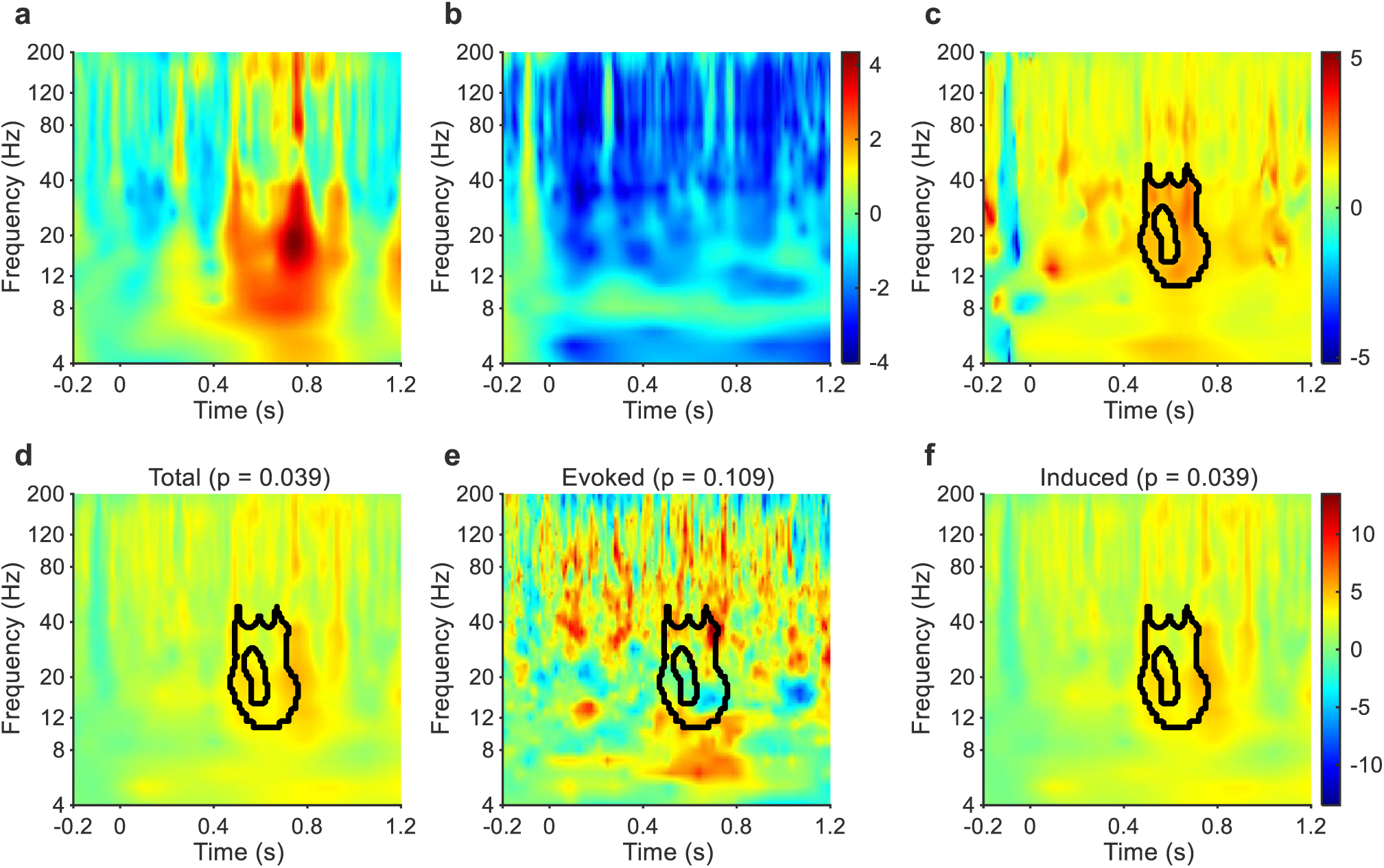
Learning-related modulation of oscillatory activity. **a, b** Time–frequency representations for Repeated (a) and Random (b) conditions. **c** Cluster-based permutation test showing increased power spanning 13–41 Hz (5th– 95th percentile range) during Repeated compared to Random trials (black contour). **d–f** Repeated minus Random difference for total (d), evoked (e), and induced (f) power, with the original cluster overlaid. (Frequency is displayed on a logarithmic scale for visualization, expanding lower frequencies to improve visualization of the beta/low-gamma cluster. Power is shown up to 200 Hz, as higher frequencies are dominated by broadband and aperiodic components; high-frequency activity was assessed separately using PAC analyses up to 550 Hz.

The cluster remained significant for total and induced (non-phase-locked) power (both p = 0.039) but not for evoked (phase-locked) power (p = 0.11). Induced activity was obtained after removal of the trial-averaged stimulus-locked response, whereas evoked activity reflected the stimulus-locked component alone. These findings indicate that the effect was driven by modulation of ongoing activity rather than consistent stimulus-evoked responses (Fig. 2d-f). No differences between conditions were observed in ongoing or prestimulus power within the cluster frequency range (both p = 0.64).

### Evolution of beta activity during learning

To assess how oscillatory activity evolved over the course of learning, we examined trial-wise power averaged across the beta–low-gamma cluster associated with the significant difference between Repeated and Random conditions. During Repeated, cluster power increased progressively within the first block and was subsequently re-established following transitions from Random to Repeated (Fig. 3a). To further characterize the progressive increase visible across Block 1, we performed a post hoc comparison of the first and final 12-trial sequence repetitions within Block 1. Cluster power was significantly greater during the final than the initial sequence repetition (p = 0.031; Fig. 3b). In contrast, no systematic change was observed in Random (p = 0.844; Fig. 3c). Together, these findings suggest that beta activity dynamically tracks the emergence and re-engagement of learned sequence structure. To assess whether learning-related changes in beta power paralleled behavioral improvement, we compared temporal changes in the group-average beta power and RT trajectories. During Repeated trials, changes in beta power showed a positive correspondence with changes in RT (r = 0.33; Fig. 3d). A circular-shift permutation test indicated that the observed alignment was greater than expected from the temporal structure of the trajectories alone, although the effect did not reach significance (permutation p = 0.062). No corresponding effect was observed during Random trials (r = 0.18, permutation p = 0.645).

**Fig. 3.**
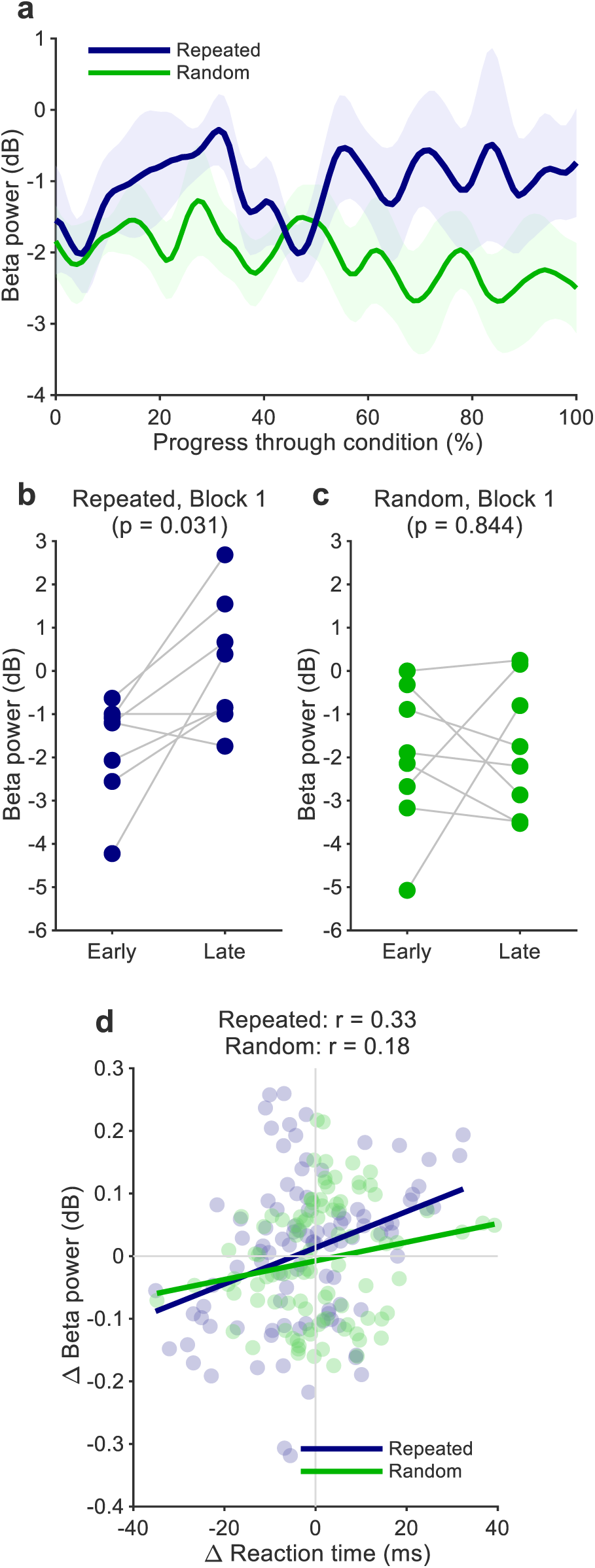
Beta dynamics during motor sequence learning. **a** Trial-wise evolution of beta power within the ERSP-defined cluster. **b**, **c** Beta power in early versus late trials within Block 1 for Repeated (b) and Random (c). Lines indicate within-subject changes. Beta power increased during Repeated (p = 0.031) but not during Random (p = 0.844). **d**. Correspondence between temporal changes in group-average reaction time (ΔRT) and beta power (Δbeta). Trajectories were temporally smoothed and transformed to first derivatives before analysis. Each point represents a normalized time point derived from the group-average trajectories. A circular-shift permutation test was used to assess whether the observed trajectory alignment exceeded that expected from the temporal structure of the signals.

### Absolute power and resting-state analyses

Absolute power within the cluster-defined frequency and time range (13–41 Hz, 500–800 ms) did not differ between Repeated and Random conditions (p = 0.58). Post-task resting recordings were available for seven participants, with 30 s analyzed per participant. Absolute power within the corresponding frequency range during post-task rest (−5.47, log₁₀-transformed absolute power) did not differ significantly from either task condition (all p ≥ 0.44). Prestimulus baseline power (−200 to 0 ms) likewise did not differ significantly between Repeated and Random trials (Repeated: −6.08; Random: −6.01; p = 0.22). Mean post-stimulus power values were similarly comparable between conditions (Repeated: −5.91; Random: −5.94). In contrast, baseline-to-post-stimulus modulation was greater during Repeated than Random trials (1.67 versus 0.67 dB). Thus, although the ERSP analysis identified significantly greater beta–low-gamma modulation during Repeated than Random trials, this effect was not accompanied by detectable differences in absolute power during the prestimulus baseline period, the post-stimulus period, or post-task rest. The observed condition difference therefore reflects differences in power modulation relative to the ongoing prestimulus task state rather than differences in absolute oscillatory power.

### Cross-frequency coupling distinguishes learning conditions

We next examined whether learning-related differences extended to cross-frequency interactions. Cluster-based permutation testing of PAC, quantified using the modulation index, revealed significant differences between the Repeated and Random conditions. A primary negative cluster (p = 0.0039, paired Cohen’s dz = 1.11) showed stronger PAC during Random than Repeated, involving beta-phase frequencies (20–30 Hz) and HFA amplitude (185– 515 Hz) (Fig. 4a-c). A smaller additional negative cluster was observed at lower beta-phase frequencies (13–19 Hz) and 185–320 Hz amplitude frequencies (p = 0.031), likewise reflecting stronger PAC during Random. These frequency ranges emerged from the data-driven cluster-based permutation analysis and were not specified a priori. Subsequent descriptive analyses focused on the primary cluster, because it showed the strongest and broadest effect. Visualization of phase-binned amplitudes derived from the primary negative cluster and of polar distributions showed similar preferred phases across conditions, with coupling peaks occurring at the rising and falling slopes of the beta cycle (Fig. 5). To quantify the apparent phase asymmetry suggested by visual inspection, HFA amplitude at phase bins closest to 90° and 270° was extracted for each participant, and a normalized asymmetry index was computed as (270° – 90°)/(270° + 90°). This revealed no significant asymmetry within either condition (Wilcoxon signed-rank tests: Repeated: p = 1.0; Random: p = 0.20) and no difference between conditions (p = 0.55). Thus, the apparent asymmetry in Random appears to reflect inter-individual variability rather than a systematic effect.

**Fig. 4.**
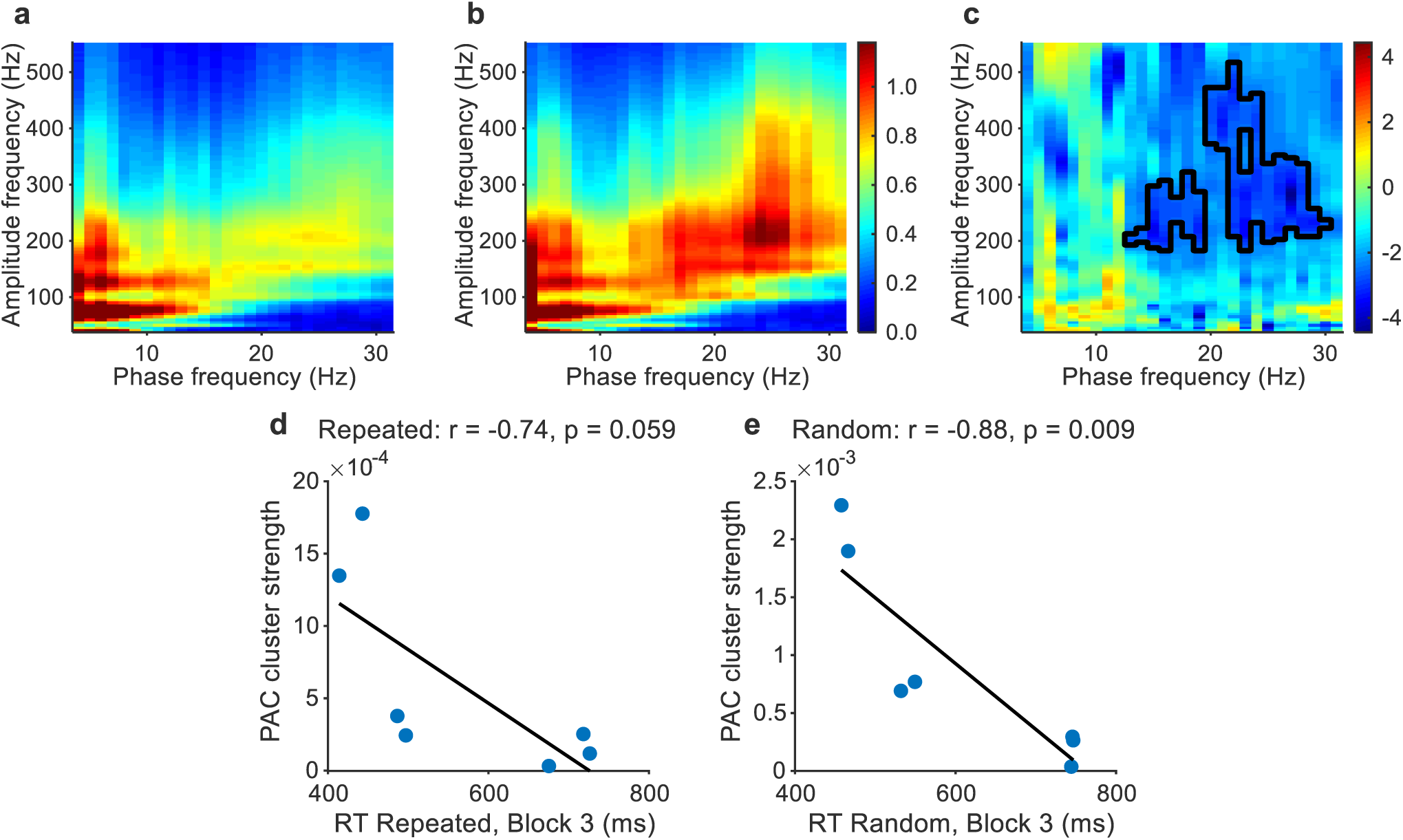
Cross-frequency coupling distinguishes sequence conditions. **a**, **b** Phase–amplitude coupling (PAC) comodulogram for Repeated (a) and Random (b). **c** Cluster-based permutation testing showing stronger beta–high-frequency activity coupling during Random than Repeated. Black outlines indicate negative clusters. **d**, **e** Relationship between reaction time (RT) and PAC cluster strength during Repeated (d) and Random (e). Lines indicate least-squares linear fits.

**Fig. 5.**
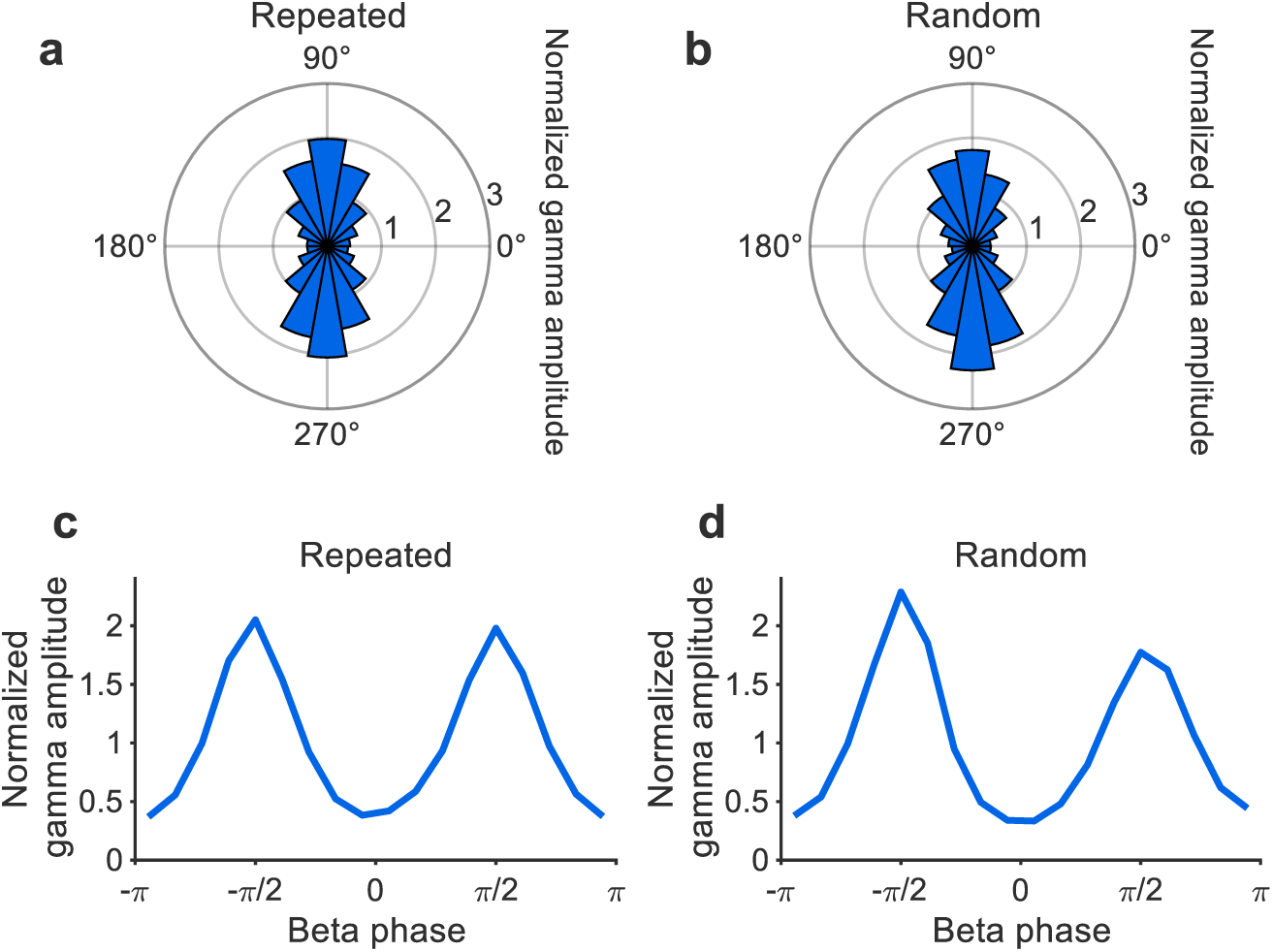
Phase-dependent modulation of high-frequency activity. **a**, **b** Phase-binned high-frequency amplitude as a function of beta phase for Repeated (a) and Random (b). **c, d** Corresponding polar histograms showing normalized amplitude across phase bins. Coupling peaked on the rising and falling slopes of the beta cycle.

Within conditions, stronger PAC tended to be associated with faster RTs. This relationship was significant in the random condition (r = −0.88, p = 0.0089) and approached significance in the repeated condition (r = −0.74, p = 0.059; Fig. 4d, e). PAC differences between conditions did not correlate with behavioral differences (r = -0.13, p = 0.79). PAC values extracted from the primary task-defined PAC range differed across rest, Repeated, and Random conditions (Friedman test: p = 0.0021). Mean PAC values were highest during Rest (0.035), lower during Random trials (0.00083), and lower still during Repeated trials (0.00056). Post hoc comparisons indicated stronger PAC during rest than during Repeated (p = 0.016) and Random trials (p = 0.031). Consistent with the primary task analysis, PAC was lower during Repeated than Random trials (p = 0.016). Thus, although the ordering of PAC values followed a pattern of Rest > Random > Repeated, the largest difference was observed between Rest and task performance, with a smaller but significant difference between Random and Repeated trials.

Together with the power analyses, the results show a striking dissociation. Oscillatory power, including in the beta range increases over the course of learning whereas beta–HFA coupling is stronger during uncertain behavior.

### PAC differences are not explained by spectral power

To ensure that the modulating oscillation in the PAC shows a peak in the power spectrum^26,27^, beta peaks were parameterized relative to the aperiodic signal component using the “fitting oscillations & one-over f” (FOOOF; f = frequency) algorithm^28,29^. Spectral parameterization identified beta peaks (13–32 Hz) in 7 of 8 participants individually in both conditions. During Repeated, the mean peak frequency was 20.6 ± 5.4 Hz and the mean peak amplitude above the aperiodic background was 0.27 ± 0.16. During Random, the mean peak frequency was 23.0 ± 5.0 Hz and the mean peak amplitude was 0.25 ± 0.13. The center frequency did not differ significantly between conditions (t(6) = -1.74, p = 0.13, Cohen’s d = -0.45) nor did peak amplitude (t(6) = 0.77, p = 0.47, Cohen’s d = 0.23). Model fits were high across participants (Repeated: R² = 0.97 ± 0.014, mean squared error [mse] = 0.018 ± 0.0094; Random: R² = 0.95 ± 0.051, mse = 0.034 ± 0.045), indicating robust separation of periodic and aperiodic components^29^.

As an additional control, extrema sharpness ratios were computed for the beta oscillation to assess whether non-sinusoidal waveform shape could account for the observed PAC. Ratios were close to 1 in both conditions (Repeated: mean ± SD: 1.0056 ± 0.0029, range: 1.0011–1.0093; Random: 1.0042 ± 0.0017, range: 1.0012–1.0061), indicating minimal waveform asymmetry and supporting the interpretation that the observed PAC reflects genuine phase-dependent modulation rather than a waveform artefact.

## Discussion

The present findings identify complementary thalamic mechanisms associated with repeated sequence execution and random trial performance. Whereas beta–low-gamma activity increased during learning compared to prestimulus baseline and tracked learned sequence structure, beta–HFA coupling was stronger during random trials and, in both conditions, associated with faster reaction times. These results suggest that the human VIM supports motor sequence learning through dissociable oscillatory mechanisms, with increased beta–low-gamma power accompanying the emergence of learned sequential structure and reduced beta–high-frequency coupling characterizing increasingly predictable task performance.

Our analysis of beta-low-gamma power during the Repeated and Random conditions was in comparison to a prestimulus baseline. In this contrast, the learning-related increase in beta–low-gamma power was driven primarily by induced rather than evoked activity, indicating modulation of ongoing oscillatory dynamics rather than stimulus-locked responses. Importantly, the prestimulus period during task performance was used as the ERSP baseline rather than a resting-state baseline. Participants were, therefore, already engaged in the task and preparing upcoming responses during the baseline interval. Additional analyses showed that absolute power within the cluster-defined frequency range did not differ significantly between Repeated, Random, and post-task resting-state recordings. Likewise, neither prestimulus nor absolute post-stimulus power differed significantly between conditions. However, ERSP measures changes in power relative to the ongoing prestimulus state within each condition rather than absolute power itself. Thus, two conditions may exhibit similar absolute power values while differing in the degree to which activity is modulated relative to the baseline. The present findings indicate that Repeated and Random trials differed primarily in how beta–low-gamma activity changed from the prestimulus period, rather than in the absolute level of oscillatory power itself. Together, these findings suggest that the ERSP effect reflects learning-related modulation relative to an already task-engaged neural state rather than a simple difference in absolute oscillatory power, whereas the PAC findings indicate that coupling between beta phase and local HFA varies across broader network states extending from rest to task performance.

Post hoc examination of cluster-averaged values indicated that power within the significant cluster was elevated relative to baseline during Repeated sequences, whereas Random trials did not differ significantly from baseline. This pattern is consistent with the interpretation that sequence learning was associated with the emergence of beta–low-gamma activity rather than simply a reduction in suppression. Increased beta power during repeated sequences is consistent with prior work linking beta oscillations to stabilized or familiar motor states^10,13,30^. In motor cortex, training-related increases in beta activity have similarly been associated with retention during motor sequence learning^30^. The temporal evolution of beta power also broadly paralleled the evolution of behavioral performance during repeated sequence execution, suggesting that learning-related beta dynamics may track the emergence of stable sequence performance at the group level. However, this correspondence, which showed a trend in the circular-shift permutation analysis, was observed at the level of group-average learning trajectories and should not be interpreted as evidence for a consistent participant-level relationship between beta power and behavioral improvement.

Temporal beta-band dynamics are similar across cortical, thalamic, and basal ganglia recordings, and both sequence learning and stimulus–response mapping are associated with modulation of beta oscillations across distributed motor networks^1,8,9,31^. Beta suppression is stronger during random than repeated sequence trials over the motor cortex^32^, whereas prolonged post-stimulus beta suppression has been associated with impaired motor sequence learning in Parkinson’s disease and essential tremor^8,33^. Conversely, VIM-DBS reduces beta power suppression at the end of sequence training, correlating with improved performance^8^. Together, these findings suggest that beta dynamics may indicate changes within the motor system related to learning. Transient beta suppression during motor task engagement is associated with a state in which the system is biased towards behavioral flexibility^34^, a state suited for learning; for example, in the Random condition, the system is challenged to identify information that might be predictive of the upcoming stimuli. The subsequent increase in beta activity has been interpreted as reflecting stabilization or maintenance of the current motor state as predictable stimulus–response relationships become established^10^.

The temporal evolution of beta activity revealed a two-stage pattern. During the initial exposure to the sequence, beta power increased progressively, whereas during subsequent repetitions, elevated beta activity was re-engaged following transitions from random to repeated sequence trials. This pattern suggests that beta oscillations track the emergence and re-expression of sequence-related structure. Consistent with this interpretation, the temporal evolution of beta power broadly paralleled the evolution of reaction time, particularly during repeated sequence execution. Notably, the significant cluster occurred relatively late after stimulus onset (approximately 500–800 ms), suggesting that it reflects learning-related modulation of beta activity during ongoing task performance rather than the classical movement-related beta decrease observed around movement initiation. More broadly, the results are consistent with proposals that beta oscillations reflect the maintenance of sensorimotor states^10^ and with evidence that the degree of beta modulation varies with motor learning^9^ and motor task demands within the VIM^13^. In a related manner, beta activity may be negatively correlated with the degree of uncertainty, or entropy of the current behavioral state, with lower beta power in the random condition when uncertainty is highest. From this perspective, the high beta observed during rest would correspond to a motor context of low uncertainty/entropy; the system is maintaining the status quo^9,10^.

In addition to changes in oscillatory power, the presence or absence of a repeated sequence modulated cross-frequency interactions: beta–HFA coupling was stronger during random trials than repeated sequences. This difference occurred in the absence of a shift in preferred phase, suggesting that the temporal organization of coupling was preserved while its strength was modulated. Increased coupling during random trials may reflect the greater computational demands associated with an uncertain behavioral context, consistent with observations of stronger coupling under conditions requiring increased cognitive or sensorimotor processing^14,35^. Notably, within each condition, higher PAC strength was associated with faster responses suggesting that cross-frequency coupling indexed the efficiency of neural processing. The absence of a relationship between condition differences in PAC and behavioral differences further supports a dissociation between mechanisms reflecting specific task demands and those supporting performance efficiency. Task-modulated cross-frequency coupling has previously been observed between low frequency VIM oscillations and motor cortical gamma activity, with thalamo–cortical PAC proposed to provide a gating mechanism for movement execution mediated by the VIM^24^. More broadly, increasing evidence suggests that the thalamus modulates information content in addition to relaying information to the cortex^15,24,35^. Within this framework, the present findings suggest that beta–HFA coupling in the VIM may reflect dynamic changes in network state associated with behavioral context and predictability.

The resting-state comparison suggests that the beta–HFA coupling identified during task performance may reflect a physiologically meaningful network state rather than an epiphenomenon of the task. Coupling was substantially stronger during rest than during task performance and remained significantly greater during Random than Repeated sequence execution. Thus, although PAC values followed a pattern of Rest > Random > Repeated, the dominant effect was a reduction in coupling during task engagement, with a smaller but significant reduction during repeated sequence execution. This pattern is consistent with the view that beta-phase modulation of local HFA reflects a state of network stabilization. During rest, strong coupling may reflect maintenance of a sensorimotor state. Engagement in motor behavior is associated with partial release from this state, while successful execution of a learned sequence is accompanied by a further reduction in coupling. In this framework, motor sequence learning is not characterized by the emergence of stronger beta-mediated coordination, but rather by a progressive decoupling of local neuronal activity from ongoing beta rhythmicity, potentially permitting greater flexibility and efficiency of information processing within thalamocortical motor networks. This interpretation is also consistent with the concomitant increase in beta power observed during Repeated relative to Random trials. Although beta power and beta–HFA coupling are often assumed to reflect related processes, the present findings suggest a dissociation between these measures. Learning was associated with stronger beta-band activity but weaker beta– HFA coupling, indicating that increased beta oscillatory power does not necessarily imply stronger beta-mediated modulation of local neuronal firing. Instead, elevated beta activity may coexist with reduced phase-locking of high-frequency activity, potentially reflecting a more efficient and less constrained network state.

Both HFA and spiking are known to be systematically modulated by the phase of lower-frequency oscillations^36,37^. Modulation of broadband HFA by beta phase is consistent with previous intracranial findings linking low-frequency oscillatory phase to local neuronal population activity^19,37^. In the motor system, beta oscillations are thought to reflect alternating states of inhibition and excitability within cortico–basal ganglia–thalamocortical circuits^10^. Notably, PAC peaked on the rising and falling slopes of the beta cycle rather than at its peaks or troughs, indicating that coupling was not centered on canonical phase extrema. Similar offsets from canonical phase extrema have been reported in previous studies of cross-frequency coupling^37,38^. The functional significance of this phase preference remains unclear, but it suggests that coupling may not be restricted to the most commonly examined phases of the beta cycle.

The presence of a spectral peak at the frequency of the modulatory oscillation is considered a prerequisite for physiologically meaningful PAC^26,27^. Spectral parameterization confirmed robust separation of periodic and aperiodic components and showed that beta peaks were present in the majority of participants, without differences in peak frequency between conditions. Thus, PAC differences emerged despite comparable beta peak characteristics across conditions. Because non-sinusoidal oscillations can produce spurious cross-frequency coupling^26^, waveform asymmetry was additionally quantified using extrema sharpness metrics^39^. Values were close to 1 in both conditions, indicating near-sinusoidal oscillations. Moreover, coupling peaked on the slopes rather than the extrema of the beta cycle. Together, these analyses support the interpretation that the observed modulation of beta–HFA coupling reflects genuine cross-frequency interactions rather than differences in underlying signal structure. Although these controls reduce the likelihood that the observed PAC reflects signal artefacts, they do not establish causality. Importantly, PAC was strongest in the condition showing lower beta–low-gamma power, arguing against a simple dependence of coupling strength on oscillatory power.

Several limitations should be considered. The sample size was constrained by the availability of intraoperative recordings. Post-task resting recordings were available for seven participants and were necessarily constrained by intraoperative time limitations. Recordings were restricted to a single effector, limiting generalizability across motor outputs. Although the study was conducted in a clinical population, the consistency of effects across participants, convergence across analyses, and clear behavioral evidence of sequence learning support the interpretation that these findings reflect general principles of thalamic network dynamics rather than essential tremor pathophysiology. Consistent with this interpretation, no differences in power or PAC were observed within the classical tremor frequency range. Finally, because the present findings are correlational, future studies combining causal manipulation of oscillatory activity with behavioral measures will be required to determine whether these dynamics directly contribute to motor sequence learning. Although the effects were robust across participants, replication in larger cohorts will be important for determining the generalizability of these findings.

The present findings reveal a dissociation between oscillatory power and cross-frequency coupling within the human VIM. Beta power increased during repeated sequence execution, whereas beta–HFA coupling was strongest in the Random condition, consistent with greater processing demands in a less predictable behavioral context. Together with the resting-state findings, these results suggest that oscillatory power and cross-frequency coupling index distinct aspects of network function. Within the cerebello–thalamo–cortical circuit, this dissociation may reflect a transition from computationally demanding processing toward more efficient and stabilized network states as learning progresses. Increasing evidence suggests that the thalamus modulates information content in addition to relaying sensory information to the cortex^15,27,40–42^. Rather than acting solely as a relay, the VIM appears to shape information flow within motor networks by modulating both the strength and timing of neural activity according to task demands. Our findings suggest that the VIM contributes not only to movement execution, but also the processing of sequential motor structure.

In conclusion, we provide evidence that the VIM is actively engaged in motor sequence learning, exhibiting distinct patterns of oscillatory activity depending on sequence structure. These findings suggest that human VIM contributes to motor learning by dynamically coordinating oscillatory states associated with learned sequential structure and less predictable behavioral contexts.

## Methods

### Participants

Eight patients with essential tremor were recruited through the Department of Stereotactic Neurosurgery, University Hospital, Magdeburg, and were studied during VIM electrode implantation or impulse-generator replacement surgery. All participants provided written informed consent prior to inclusion. The study was approved by the local ethics committee of the Medical Faculty of the Otto von Guericke University Magdeburg and carried out in accordance with the Declaration of Helsinki.

### Study design

Each participant completed one session of the SRTT^6,8,9,25,33^, implemented using Presentation software (Neurobehavioral Systems, Berkeley, CA, USA). Participants viewed four empty black square outlines on a gray background, arranged horizontally on a computer screen. When a square was filled in red, the participant pressed the corresponding button on an ergonomically shaped response pad, as quickly and accurately as possible, using the four fingers of the left hand. The stimuli followed either a fixed 12-item sequence (1-3-2-1-4-1-2-3-1-3-2-4) or a pseudorandom order. In the random condition, each location was highlighted at least once within every 12 trials, and immediate repetitions were not permitted. Stimuli were presented for 500 ms, with a fixed stimulus onset asynchrony of 1200 ms, independent of response time. The experiment comprised three blocks, separated by one-minute breaks. Within each block, the 12-item sequence was repeated three times (36 trials), followed by 36 random trials, a further three sequence repetitions, and a final 36 random trials, yielding 144 trials per block.

Post-task resting data were recorded directly after completion of the experiment, when intra-operative time permitted, following the final task epoch. Participants were included when at least 30 s of post-task data were available. To minimize contamination by immediate post-task movement or settling, the final 30 s of the available post-task segment were extracted for each included participant. Resting data meeting these criteria were available for 7 of the 8 participants included in the electrophysiological analyses.

### Intracranial recording and preprocessing

Local field potentials were recorded from implanted DBS electrodes in the right VIM. Electrodes comprised either four ring contacts (Medtronic, Dublin, Ireland) or segmented leads (Abbott, North Chicago, IL, USA); segmented contacts were averaged within each level. Signals were referenced to the electrode casing and grounded via a scalp electrode over the left parietal cortical region (Brain Products, Gilching, Germany). Data were sampled at 20 kHz and recorded using Inomed software (Inomed, Emmendingen, Germany). Data were processed offline in MATLAB R2023b/2025a using Fieldtrip (version 20240614)^43^ and Brainstorm^29^. Signals were visually inspected for artefacts, and no additional rejection procedures were applied to preserve trial counts. Signals were re-referenced bipolarly by subtracting adjacent contacts to attenuate noise, increase spatial selectivity, and reduce effects of volume conduction from distant sources^44–46^, yielding three adjacent bipolar channels per participant from the four contact-level channels.

Structural magnetic resonance images were recorded pre-operatively using a Siemens Verio scanner (Siemens, Erlangen, Germany) with a 32-channel head coil. VIM-DBS electrode coordinates were established through co-registration of these images with post-operative computed tomography, combined with intraoperative stereotactic x-ray coordinates, and are reported in relation to the anterior–posterior commissural line. The center coordinates of the VIM were determined by the neurosurgeon using the Inomed software package. For further analysis, we selected the electrode closest to the VIM center for each participant.

### Time–frequency analysis

Continuous data were notch-filtered at 47–53 Hz and its harmonics. Data were epoched from -1400 ms to 2600 ms relative to stimulus onset. Time–frequency decomposition was performed using six-cycle complex Morlet wavelets (4–200 Hz in 1 Hz steps; 10 ms resolution) over the interval -200 ms to 1200 ms. Power was normalized to a prestimulus baseline window (-200 to 0 ms) using a decibel transform (10·log10(power/baseline power)). ERSPs were computed for each participant by averaging trials separately for repeated and random conditions, and grand averages were calculated across participants for visualization. Trials with incorrect or missing responses were excluded from ERSP analyses. Condition differences were assessed using cluster-based permutation tests. To characterize the spectral extent of significant clusters, the frequency distribution of cluster samples was extracted, and the 5th–95th percentile range was calculated. Significant clusters were subsequently used to define regions of interest for follow-up analyses.

To further characterize the direction of the significant ERSP effect, mean power values were extracted separately for each participant and condition from the frequency and time range corresponding to the 5th–95th percentile boundaries of the significant cluster (13–41 Hz, 500–800 ms). Power was averaged across all frequency–time samples within this cluster-defined range. Because ERSP values were expressed relative to the prestimulus baseline using a decibel transform, a value of 0 dB corresponds to baseline power. Mean values for Repeated and Random conditions were compared against zero using two-tailed Wilcoxon signed-rank tests.

#### Induced vs. evoked activity

To determine whether the observed time–frequency effect reflected evoked (phase-locked) or induced (non-phase-locked) activity^47,48^, the signals were decomposed into total, evoked, and induced components. For each participant and condition, time–frequency decomposition was applied to single-trial data using wavelet convolution, with identical time–frequency parameters and baseline normalization procedures to those in the primary ERSP analysis. Evoked activity was estimated by first averaging trials in the time domain to obtain the event-related potential (ERP) and then performing the time–frequency decomposition of this averaged signal. Induced activity was obtained by subtracting the ERP from each individual trial prior to time–frequency analysis, thereby isolating non-phase-locked oscillatory activity. The resulting time–frequency representations were subsequently normalized using the same prestimulus baseline (−200 to 0 ms) as the primary ERSP analysis. Differences between conditions for total, evoked, and induced power were assessed using paired Wilcoxon signed-rank tests.

#### Baseline changes vs. stimulus-related dynamics

To assess whether the ERSP cluster reflected baseline oscillatory activity changes or stimulus-related dynamics, ongoing power outside the peri-stimulus interval (0 to 1200 ms), prestimulus power (-500 to 0 ms), and induced ERSP cluster power were computed. Ongoing and prestimulus power were quantified within the frequency range of the ERSP cluster, while induced power was extracted from the exact cluster mask. Condition differences were assessed using paired Wilcoxon signed-rank tests.

#### Time-resolved beta power analysis

To examine slow temporal dynamics of beta-band activity across the course of each condition, trial-wise estimates of beta power were computed for each participant by averaging power within the frequency–time region defined by the significant ERSP cluster. For each condition, these trial-wise time series were smoothed using a Gaussian kernel (window length: 24 trials) to reduce high-frequency trial-to-trial variability while preserving slower fluctuations in oscillatory activity.

#### Initial learning

Beta power within the ERSP cluster was averaged across trials belonging to the first and last 12-item sequences of Block 1 (early vs. late) for repeated sequence and random trials. Differences were assessed using two-tailed Wilcoxon signed-rank tests across participants.

#### Relationship between beta and behavioral learning trajectories

To assess whether neural and behavioral dynamics co-evolved over the course of the task, we quantified the correspondence between changes in beta power (Δbeta power) and changes in reaction time (ΔRT). Group-level beta-power trajectories were derived from trial-wise ERSP cluster values, and RT trajectories were derived from sequence-averaged behavioral data. Beta trajectories were smoothed using a Gaussian kernel of 24 trials and RT trajectories using a Gaussian kernel of two sequence bins. Both measures were interpolated to a common normalized time axis (0–100% of condition duration) to account for differences in trial counts across participants. Change scores were computed as the first temporal derivative of the group-averaged trajectories, yielding Δbeta power and ΔRT. Pearson’s correlation coefficient was used to quantify the correspondence between Δbeta power and ΔRT separately for repeated and random conditions. Because these analyses were based on temporally smoothed group-level trajectories, statistical significance was assessed using a circular-shift permutation test rather than the analytical p-value of the correlation coefficient. One trajectory was circularly shifted relative to the other by a random offset while preserving its temporal structure, and the correlation was recalculated. This procedure was repeated 10,000 times to generate a null distribution. The empirical two-sided p-value was calculated as the proportion of permuted correlations whose absolute value equaled or exceeded that of the observed correlation. Scatter plots with least-squares regression lines are shown for visualization only.

#### Absolute power and resting-state analyses

To assess whether the observed ERSP effect reflected differences in absolute oscillatory power, additional analyses were performed on non-baseline-corrected data. Stimulus-locked epochs (−1.4 to 2.6 s relative to stimulus onset) were extracted using the same preprocessing pipeline as the ERSP analysis. Time–frequency representations were computed using Morlet wavelets (6 cycles) without baseline normalization. Mean absolute power was then extracted from the frequency and time range corresponding to the ERSP cluster (13–41 Hz, 500–800 ms). Power was compared between Repeated and Random conditions using Wilcoxon signed-rank tests.

To further assess the relationship between task-related and resting activity, the same frequency range (13–41 Hz) was quantified during post-task resting recordings. Resting-state power was computed from the final 30 s of the available post-task segment and compared with Repeated and Random task activity using Friedman tests with Wilcoxon signed-rank post hoc comparisons. In addition, baseline (−200 to 0 ms), post-stimulus (500–800 ms), and baseline-normalized (dB) power were compared between conditions within the cluster-defined frequency range.

### Phase–amplitude coupling

For the PAC analysis, data were segmented into 12-item sequence epochs (14.4 s), stimulus-locked to the first trial in each sequence. Longer data segments were required to ensure reliable estimation of coupling strength^49^. Phase and amplitude were extracted using complex Morlet wavelets (phase: 4-31 Hz in 1 Hz steps and 10 ms resolution, six-cycle wavelets; amplitude: 40-550 Hz in 5 Hz steps and 10 ms resolution, seven-cycle wavelets), yielding phase and amplitude estimates comparable to Hilbert-based approaches^50^. Phase and amplitude extraction using complex Morlet wavelets with six cycles for phase frequencies and seven cycles for amplitude frequencies followed prior wavelet-based PAC approaches using frequency-range-specific cycle settings^51^. PAC was calculated for each trial using the modulation index^38^ and averaged within participants. Cluster-based permutation testing was applied, and subsequent analyses focused on cluster-defined frequency ranges.

To further characterize the significant PAC effect, mean modulation index values were extracted for each participant and condition from the phase and amplitude frequency ranges corresponding to the 5th–95th percentile boundaries of the primary significant PAC cluster (20–30 Hz phase, 185–515 Hz amplitude). Modulation index values were averaged across all phase–amplitude frequency pairs within this cluster-defined range. Values were compared across post-task Rest, Repeated, and Random conditions using Friedman tests with Wilcoxon signed-rank post hoc comparisons. For the resting-state analyses, the final 30 s of post-task rest were divided into two non-overlapping 14.4 s epochs to match the epoch duration used for task PAC estimation.

To verify the presence of oscillatory peaks in the modulating frequency range^26,27^, power spectral density was parameterized using the “fitting oscillations and one-over f” (FOOOF; f = frequency) algorithm, as implemented in the Brainstorm toolbox, to separate periodic and aperiodic components^28,29^. Model fit quality was assessed using the coefficient of determination (R²) and mean squared error (MSE), and summarized across participants (mean ± SD), following standard practice in spectral parameterization^28^.

To control for potential confounds from non-sinusoidal waveform shape of the modulating oscillation^26^, extrema sharpness ratios were computed^39^. Peaks and troughs were identified in the beta-filtered signal, and sharpness was quantified for each extremum as the difference between the signal value at the peak (or trough) and the mean of the surrounding points within a small temporal window (±5 ms). Mean sharpness values were computed separately for peaks and troughs, and the extrema sharpness ratio was defined as the ratio of the larger to the smaller value. Values close to 1 indicate a near-sinusoidal waveform, whereas larger values indicate increasing asymmetry. Summary statistics were computed across participants.

To visualize the preferred phase of HFA, phase–amplitude distributions were computed. The frequency ranges for phase and amplitude were derived from the significant cluster identified in the cluster-based permutation test of PAC. The signals were band-pass filtered using zero-phase finite impulse response filters in two frequency ranges: a low-frequency band providing the phase (beta range) and a high-frequency band providing the amplitude envelope. The instantaneous phase of the low-frequency signal was obtained using the Hilbert transform. The amplitude envelope of the high-frequency signal was computed as the absolute value of the analytic signal obtained via the Hilbert transform. To avoid edge artefacts, data at the beginning and end of the recording (1 s) were discarded. HFA amplitude was averaged within 18 phase bins (−π to π), normalized within participants, and then averaged across participants. Distributions were visualized in polar coordinates, where the radius represents normalized HFA amplitude and the angle represents the phase of the low-frequency oscillation.

#### Cross-frequency coupling and behavior

To assess whether cross-frequency coupling was related to behavioral performance, mean PAC strength was extracted for each participant from the significant cluster identified in the group-level analysis. PAC values were averaged across the cluster mask separately for Repeated and Random, yielding a single value per participant and condition. Behavioral performance was quantified as the mean RT within the third (final) block of each condition. Correlations between PAC strength and RTs were computed separately for Repeated and Random using Pearson’s correlation coefficient. In addition, correlations between condition differences in PAC (Random minus Repeated) and RT (Random minus Repeated) were assessed.

### Statistics

#### Behavioral analysis

Statistical analysis was performed using IBM SPSS Statistics 23 (IBM, Armonk, NY, USA), and visualized with MATLAB 2025a (MathWorks, Natick, MA, USA). RTs were analyzed with a repeated measures ANOVA with the within-subject factors *Time* (Blocks 1 to 3) and *Sequence type* (Repeated, Random). Post hoc tests are reported following Bonferroni correction for multiple comparisons. One participant completed only the first block of the experiment and was therefore excluded from the behavioral ANOVA. Electrophysiological data from the completed block were retained for analyses whenever possible.

#### Electrophysiological analysis

For the time–frequency analysis, non-parametric cluster-based permutation testing^52^ was performed across all time points (−200 to 1200 ms) and frequencies (4–200 Hz) to compare ERSPs between repeated sequences and random trials. A similar procedure was applied to PAC, using phase frequencies from 4 to 31 Hz and amplitude frequencies from 40 to 550 Hz. Cluster formation was based on dependent-samples two-sided t-tests with a cluster-forming threshold of p < 0.025. Condition labels were permuted within participants by randomly exchanging repeated sequence and random trials, implemented as within-subject sign-flipping. Given the sample size, all possible permutations (2⁸ = 256) were evaluated, resulting in an exact permutation test. For each permutation, the maximum cluster-level statistic (sum of t-values within each cluster) was retained to construct the null distribution. The observed cluster statistics were compared against this distribution, and p-values were determined. Effect sizes for significant clusters were calculated using Cohen’s d_z_, defined as the mean of the paired differences divided by their standard deviation.

## Data availability statement

Raw intracranial electrophysiological data are not publicly available because of participant privacy and ethical restrictions but are available from the corresponding author upon reasonable request and subject to institutional data-sharing agreements. Source data underlying the graphs and statistical analyses will be deposited in a public repository and made available upon publication.

## Code availability statement

Custom MATLAB scripts used for the analyses and figure generation will be deposited in a public repository and made available upon publication. FieldTrip^43^ (v20240614) and Brainstorm^29^ were used for electrophysiological analyses.

## Acknowledgements

This work was supported by the Deutsche Forschungsgemeinschaft (DFG) [grant number SW 214/2-1 (C.M.S.-R.)].

## Author contributions

Conceptualization, methodology, software, formal analysis (behavioral and electrophysiological analyses, including ERSP, PAC, resting-state, and extended analyses for the final manuscript), visualization, writing – original draft, supervision, and funding acquisition: C.M.S.-R. Patient recruitment: L.B. and C.M.S.-R. Investigation (intraoperative data collection): C.M.S.-R., L.B., S.R., and M.D. Data preprocessing: A.V. Formal analysis (initial behavioral, ERSP, and PAC analyses): A.V. Interpretation: A.V., S.J.N., A.d.O.A., M.D., R.T.K., R.B.I., and C.M.S.-R. Writing – review and editing: A.V., L.B., S.J.N., A.d.O.A., M.D., R.T.K., R.B.I., and C.M.S.-R. All authors read and approved the final manuscript.

## IRB statement

The Local Ethics Committee of the University Hospital Magdeburg granted ethical approval (134/18). All participants provided informed, written consent before study inclusion, in accordance with the Declaration of Helsinki, and were informed of their right to cease participation at any time without providing reasons.

## References

1. Klostermann, F. et al. Task-related differential dynamics of EEG alpha- and beta-band synchronization in cortico-basal motor structures. Eur. J Neurosci. 25, 1604–15 (2007).

2. Hardwick, R. M., Rottschy, C., Miall, R. C. & Eickhoff, S. B. A quantitative meta-analysis and review of motor learning in the human brain. Neuroimage 67, 283–97 (2013).

3. Chen, H., Hua, S. E., Smith, M. A. & Lenz, F. A. Effects of human cerebellar thalamus disruption on adaptive control of reaching. Cereb. Cortex 16, 1462–73 (2006).

4. Benabid, A. L. et al. Long-term suppression of tremor by chronic stimulation of the ventral intermediate thalamic nucleus. The Lancet 337, 403–6 (1991).

5. Cury, R. G. et al. Thalamic deep brain stimulation for tremor in Parkinson disease, essential tremor, and dystonia. Neurology 89, 1–8 (2017).

6. Terzic, L. et al. Deep brain stimulation of the ventrointermediate nucleus of the thalamus to treat essential tremor improves motor sequence learning. Hum. Brain Mapp. 43, 4791–9 (2022).

7. Tan, H. et al. Decoding voluntary movements and postural tremor based on thalamic LFPs as a basis for closed-loop stimulation for essential tremor. Brain Stimul. 12, 858–867 (2019).

8. Voegtle, A. et al. Ventrointermediate thalamic stimulation improves motor learning in humans. Commun. Biol. 7, (2024).

9. Pollok, B., Latz, D., Krause, V., Butz, M. & Schnitzler, A. Changes of motor-cortical oscillations associated with motor learning. Neuroscience 275, 47–53 (2014).

10. Engel, A. K. & Fries, P. Beta-band oscillations-signalling the status quo? Curr. Opin. Neurobiol. 20, 156–165 (2010).

11. Basha, D. et al. Beta oscillatory neurons in the motor thalamus of movement disorder and pain patients. Exp. Neurol. 261, 782–790 (2014).

12. Kilavik, B. E., Zaepffel, M., Brovelli, A., MacKay, W. A. & Riehle, A. The ups and downs of beta oscillations in sensorimotor cortex. Exp. Neurol. 245, 15–26 (2013).

13. Basha, D. et al. Beta band oscillations in the motor thalamus are modulated by visuomotor coordination in essential tremor patients. Front. Hum. Neurosci. 17, 1082196 (2023).

14. Canolty, R. T. & Knight, R. T. The functional role of cross-frequency coupling. Trends Cogn. Sci. 14, 506–515 (2010).

15. Sweeney-Reed, C. M. et al. Corticothalamic phase synchrony and cross-frequency coupling predict human memory formation. Elife 3, e05352 (2014).

16. Florin, E. & Baillet, S. The brain’s resting-state activity is shaped by synchronized cross-frequency coupling of neural oscillations. Neuroimage 111, 26–35 (2015).

17. Lisman, J. E. & Jensen, O. The theta-gamma neural code. Neuron 77, 1002–16 (2013).

18. Combrisson, E. et al. From intentions to actions: Neural oscillations encode motor processes through phase, amplitude and phase-amplitude coupling. Neuroimage 147, 473–487 (2017).

19. De Hemptinne, C. et al. Exaggerated phase-amplitude coupling in the primary motor cortex in Parkinson disease. Proc. Natl. Acad. Sci. U. S. A. 110, 4780–4785 (2013).

20. Yanagisawa, T. et al. Regulation of motor representation by phase–amplitude coupling in the sensorimotor cortex. J Neurosci 32, 15467–75 (2012).

21. De Hemptinne, C. et al. Therapeutic deep brain stimulation reduces cortical phase-amplitude coupling in Parkinson’s disease. Nat. Neurosci. 18, 779–786 (2015).

22. Gong, R. et al. Spatiotemporal features of β-γ phase-amplitude coupling in Parkinson’s disease derived from scalp EEG. Brain 144, 487–503 (2021).

23. Schnitzler, S. et al. Occurrence of thalamic high frequency oscillations in patients with different tremor syndromes. Clin. Neurophysiol. 129, 959–966 (2018).

24. Opri, E., Cernera, S., Okun, M. S., Foote, K. D. & Gunduz, A. The functional role of thalamocortical coupling in the human motor network. J. Neurosci. 39, 8124–8134 (2019).

25. Nissen, M. & Bullemer, P. Attentional requirements of learning: evidence from performance measures. Cogn. Psychol. 19, 1–32 (1987).

26. Aru, J. et al. Untangling cross-frequency coupling in neuroscience. Curr. Opin. Neurobiol. 31, 51–61 (2015).

27. Sweeney-Reed, C. et al. Anterior thalamic high frequency band activity is coupled with theta oscillations at rest. Front. Hum. Neurosci. 11, 358 (2017).

28. Donoghue, T. et al. Parameterizing neural power spectra into periodic and aperiodic components. Nat. Neurosci. 23, 1655–1665 (2020).

29. Tadel, F., Baillet, S., Mosher, J. C., Pantazis, D. & Leahy, R. M. Brainstorm: a user-friendly application for MEG/EEG analysis. Computational Intell. and Neurosci. 2011, (2011).

30. Dyck, S. & Klaes, C. Training-related changes in neural beta oscillations associated with implicit and explicit motor sequence learning. Sci. Rep. 14, (2024).

31. Lum, J. A. G. et al. Neural basis of implicit motor sequence learning: modulation of cortical power. Psychophysiology 60, e14179 (2022).

32. Voegtle, A. et al. Cholinergic modulation of motor sequence learning. Eur. J. Neurosci. 60, 3706–18 (2024).

33. Meissner, S. N., Krause, V., Südmeyer, M., Hartmann, C. J. & Pollok, B. Pre-stimulus beta power modulation during motor sequence learning is reduced in Parkinson’s disease. NeuroImage: Clin. 24, 102057 (2019).

34. Alegre, M. et al. Alpha and beta changes in cortical oscillatory activity in a go/no go randomly-delayed-response choice reaction time paradigm. Clin. Neurophys. 117, 16–25 (2006).

35. Jensen, O., Gips, B., Bergmann, T. O. & Bonnefond, M. Temporal coding organized by coupled alpha and gamma oscillations prioritize visual processing. Trends Neurosci. 37, 357–369 (2014).

36. Meij, R. Van Der, Kahana, M. & Maris, E. Phase–amplitude coupling in human electrocorticography Is spatially distributed and phase diverse. J. Neurosci. 32, 111–23 (2012).

37. Canolty, R. T. et al. High gamma power is phase-locked to theta oscillations in human neocortex. Science (1979). 313, 1626–8 (2006).

38. Tort, A. B. L., Komorowski, R., Eichenbaum, H. & Kopell, N. Measuring phase-amplitude coupling between neuronal oscillations of different frequencies. J. Neurophysiol. 104, 1195–1210 (2010).

39. Cole, S. R. & Voytek, B. Brain oscillations and the importance of waveform shape. Trends Cogn. Sci. 21, 137–49 (2017).

40. Murray Sherman, S. The thalamus is more than just a relay. Curr. Biol. 17, 417–22 (2007).

41. Sweeney-Reed, C. M. et al. The role of the anterior nuclei of the thalamus in human memory processing. Neurosci. Biobehav. Rev. 126, 146–58 (2021).

42. Sweeney-Reed, C. & Knight, R. Memory and the human anterior thalamus. in The Cerebral Cortex and Thalamus (eds. Usrey, W. M. & Sherman, S. M.) vol. Section VIII Chapter 43 (Oxford University Press, New York, 2023).

43. Oostenveld, R., Fries, P., Maris, E. & Schoffelen, J. M. FieldTrip: Open source software for advanced analysis of MEG, EEG, and invasive electrophysiological data. Comput. Intell. Neurosci. 2011, 156869 (2011).

44. Mercier, M. R. et al. Advances in human intracranial electroencephalography research, guidelines and good practices. Neuroimage 260, (2022).

45. Meisler, S. L., Kahana, M. J. & Ezzyat, Y. Does data cleaning improve brain state classification? J. Neurosci. Meth. 328, (2019).

46. Beck, A. K. et al. Thalamic and basal ganglia regions are involved in attentional processing of behaviorally significant events: evidence from simultaneous depth and scalp EEG. Brain Struct. Funct. 223, 461–474 (2018).

47. David, O., Kilner, J. M. & Friston, K. J. Mechanisms of evoked and induced responses in MEG/EEG. Neuroimage 31, 1580–1591 (2006).

48. Tallon-Baudry, C., Bertrand, O., Delpuech, C. & Pernier, J. Stimulus specificity of phase-locked and non-phase-locked 40 Hz visual responses in human. J. Neurosci. 16, 4240–4249 (1996).

49. Dvorak, D. & Fenton, A. A. Toward a proper estimation of phase-amplitude coupling in neural oscillations. J. Neurosci. Meth. 225, 42–56 (2014).

50. Bruns, A. Fourier-, Hilbert- and wavelet-based signal analysis: are they really different approaches? J. Neurosci. Meth. 137, 321–332 (2004).

51. Hirano, S. et al. Phase-Amplitude Coupling of the Electroencephalogram in the Auditory Cortex in Schizophrenia. Biol. Psychiatry Cogn. Neurosci. Neuroimaging 3, 69–76 (2018).

52. Maris, E. & Oostenveld, R. Nonparametric statistical testing of EEG- and MEG-data. J. Neurosci. Meth. 164, 177–90 (2007).

